# Identification and mitigation of blood’s interference with the antimicrobial activity of AgNbO_3_ particles

**DOI:** 10.1101/2024.10.19.619205

**Authors:** Cyrus Talebpour, Fereshteh Fani, Marc Ouellette, Marilynn Fairfax, Houshang Alamdari, Hossein Salimnia

## Abstract

The detrimental impact of blood on the antimicrobial activity of AgNbO_3_ particles was identified and investigated. It was observed that the impact is more severe in the case of lysed blood and also operates in the case of commonly used silver salt, AgNO_3_. The inhibition was shown to be due to hemoglobin, but unrelated to the heme moiety. In an attempt to find additives to mitigate the inhibitory effect of hemoglobin, iron ions and the chelating agent, K_2_EDTA, were selected as promising candidates. Including iron on the particles was shown to have a marginal effect, but supplying the medium with K_2_EDTA, an agent preventing clotting of blood samples, fully countered the deleterious impact of hemoglobin on AgNbO_3_ activity. These findings may be relevant for adapting the silver compounds to applications such as wound dressings, where silver’s antimicrobial action would have to take place in a medium containing blood.

## 1. Introduction

Silver compounds, including silver salt AgNO_3_, silver nanoparticles, and silver zeolites have been intensively studied as broad-spectrum and relatively safe replacements for antibiotics in diverse biomedical applications [1–4]. Although different mechanisms of action have been proposed to cause the antibacterial activity, there is general agreement on the central role of silver ions, which become available through dissociation, corrosion, or ion exchange [5]. The elution of ions from relatively stable silver nanoparticles, which occurs via the intermediate formation of Ag_2_O on the surface of the nanoparticles by dissolved oxygen or hydrogen peroxide formed as metabolic product of a nearby microorganism, is thought to have a central role in antimicrobial action [6]. Release of silver ions results in the loss of antimicrobial activity over time and may compromise biocompatibility due to potential risks when the compound is in clinical applications, such as wound-dressing or medical implants. To achieve a more biocompatible alternative we fixed the silver atoms in a stable crystal structure (AgNbO_3_) to protect it from corrosion and dissolution [7]. Since untreated AgNbO_3_ does not have noticeable antibacterial activity, we came up with a series of thermo-mechano- chemical processes to successfully activate AgNbO_3_ particles by subjecting them to harsh impact and shearing forces. The resulting nanostructured particles may be a suitable replacement for antibiotics or conventional silver in areas such as antimicrobial dental implants [8], bone cement [9], wound dressings [10], and hip and knee implant devices [11, 12], thanks to their durability, corrosion resistance, and negligible levels of toxicity to the surrounding tissue.

Most of the reported activity assays have been performed in culture media without the addition of human blood components or other biologically relevant compounds. Thus, their results are not necessarily transferrable to complex biological samples where the presence of interfering material may affect the antimicrobial activity of Ag^+^ in vivo [13, 14]. This point has been discussed by Croes et al with respect to the poor bactericidal effects of Ag-based coatings in bone infection models [15].

Since the issue of reduced activity of silver ions under physiologic conditions was recognized, strategies for addressing it were also investigated. For instance, Vasiliev et al have shown that addition of CuSO_4_, resulting in the release of copper ions, can counter the impact of serum proteins on the activity of AgNO_3_ [16]. This is an instance of synergism of silver/transition metals, which has been demonstrated to result in increases of up to 8-fold in antimicrobial activity against certain bacterial species [17, 18].

The above observations highlight the importance of identifying and countering the effects of interfering biological materials in order to establish the suitability of a target antimicrobial for a prospective application. In this study, we investigate the potential inhibitory impact of blood on the antimicrobial activity of the AgNbO_3_ particles. We first investigated this effect by monitoring the inhibition of the antimicrobial effect of the particles in the presence of whole and lysed blood. Based on the results obtained here, we next determined the adverse effects of suspected blood components. Finally, we shifted our focus to mitigating the impact of implicated blood components by selected additives, including iron ions and the chelating agent, K_2_EDTA.

## 2. Methodology

### 2.1 Preparation and characterization of nanostructured antimicrobial particles

Nanostructured AgNbO_3_ antimicrobial particles were synthesized using the Activated Reactive Synthesis (ARS) method described previously [7]. Briefly, the method consists of the mixing of Ag_2_O (Sigma-Aldrich Corp) and Nb_2_O_5_ (Inframat^Ⓡ^ Advanced Materials LLC) in a 1 g to 1.147 g weight ratio respectively; The mixture was calcined so the material undergoes the reaction necessary to form AgNbO_3_. The calcined powder was subjected to successive steps of high and low energy ball milling, which yielded nanostructured AgNbO_3_ particles. We previously reported that these particles had a specific surface area of 7.9 m^2^/g and the average particle size was 438 nm. The silver release rate of the particles was less than 1% of its total mass after 35 days of storage in water.

A second batch of nanostructured AgNbO_3_ particles with substitutional doping of Fe_2_O_3_ was synthesized using the same ARS procedure. The only difference was to mix the raw materials Ag_2_O, Nb_2_O_5_ and Fe_2_O_3_ (Sigma-Aldrich Corp) with the weight ratio of 1 g to 1.090 g to 0.0345 g respectively to achieve AgFe_0.05_Nb_0.95_O_3_.

### 2.2 Characterization of nanostructured antimicrobial particles for surface composition

The surface composition of the particles was interrogated by photoelectron spectroscopy (XPS) with an analysis depth of ∼10 nm and surface area of 100 µm × 100 µm. The analysis was carried out using the PHI Quantes X-Ray Photoelectron Spectrometer. The survey spectra and the high-resolution spectra were acquired using the monochromatized Kɑ line of standard aluminum (hv = 1486.6 eV) with charge compensation. Photoelectron detection was performed at a take-off angle of 45° with respect to the sample surface for both survey and high-resolution spectra. High resolution C1s spectra were recorded and calibrated at 285 eV for the analysis. The analysis chamber was set to vacuum pressure at 10^-10^ mbar. The analysis of the results was performed using the open source CasaXPS software and the appropriate elemental library for the XPS machine was used to apply the correct relativity sensitivity factor for each atomic level.

### 2.3 Bacterial strain and culture conditions

The bacterial strain used in this study was *Escherichia coli* ATCC # 25922. Cell stock in 50% glycerol was taken from a −80 °C freezer. After thawing, 30-50 µL was transferred to a Tryptic Soy Agar (TSA) (with 5% sheep blood) plate (the P1 plate), inoculated by standard streaking, and incubated at 37 °C overnight under aerobic condition until colonies were visible. A well-formed, representative colony from the plate was picked and inoculated into 3 mL of Tryptic Soy Broth (TSB) and incubated at 37 °C with 150 rpm shaking for 3 h. Then, 1 mL aliquots were transferred to 2 mL sterile Eppendorf tubes and centrifuged at 8000 rpm for 8 min in a microcentrifuge. The harvested cells were resuspended in 1 mL TSB and 0.5 mL was used for OD_600_ measurement. Using an in-house OD_600_ vs cell count correlation database, a cell suspension of 1.5 × 10^8^ CFU/mL was prepared. The suspension was then diluted 1/100 in TSB to a final concentration of 1.5 × 10^6^ CFU/mL.

### 2.4 Antimicrobial activity on different agar media

Three types of agar plates obtained from Becton Dickinson, including Muller Hinton Agar (MHA), Blood Agar Plate (BAP) and Chocolate Agar Plate (CAP), were used to assess the impact of their composition on the antimicrobial activity of microbial cells. First, on the back of the plate guiding circles were marked to facilitate the two approaches described below.

1. A 5 µL of aliquot of particle suspension, containing preselected concentration of particles (1600, 800, 400, 200, 100, 50 and 0 µg/mL), was dispensed at the center of the designated circles. The particle suspension spread spontaneously to a diameter of ∼8 mm and was allowed to air dry for approximately 30 min. According to our previous analysis [19], these circles contained 160, 80, 40, 10, and 5 ng/mm^2^ of particles respectively. Then, 1 µL of bacterial suspension, having a nominal concentration of 10^5^ organisms/mL was dispensed in the center of the particle spot. The plates were incubated for 6 h in 37 °C, then left at room temperature overnight, and inspected the next day for bacterial growth. The transfer to room temperature slowed bacterial replication, in order to restrict confluent growth, facilitating the ability to distinguish cases of non-inhibition from partial inhibition.
2. Bacterial cell suspension (1/100 dilution in TSB of a 0.5 McFarland) was spread over the entire surface of the plate using a swab that had been immersed in the cell suspension tube. Then 5 µL of AgNbO_3_ dilutions (50-1600 µg/ml) were loaded onto their designated locations on the plate. For a positive control, 5µL of water was dispensed in the circle at the center of the plate. The plate was incubated at 35 °C overnight and then photographed. This test was also repeated to determine the interaction with different concentrations of AgNO_3_ as a reference material.

### 2.5 Measuring the minimum inhibitory concentration by the microdilution method

The antimicrobial activity of AgNbO_3_ particles was measured by the broth microdilution method by growing bacterial cells inside a series of wells on a microwell plate containing growth media. Each well was supplied with the concentrations of the antimicrobial agent to be tested, in a series of one to two dilutions from one well to the next. Bacterial cells with the target concentration of 1.5 × 10^6^ CFU/mL were dispensed into each well. After overnight incubation, the wells were visually inspected for signs of growth. The minimum concentration of the particles for which no growth was observed was taken as the minimum inhibitory concentration (MIC) value.

### 2.6 Measuring the antimicrobial activity of AgNbO_3_ particles in the presence of interfering materials

The antimicrobial activity of AgNbO_3_ particles was measured by the broth microdilution method by growing bacterial cells in a series of wells on a microwell plate containing growth media having different levels of hemin (Sigma-Aldrich Corp), hemoglobin (Sigma-Aldrich Corp) FeSO_4_ (Sigma-Aldrich Corp) and FeCl_3_ (Sigma-Aldrich Corp). Each well was supplied with different concentrations of the antimicrobial agent, in a serial one to two dilutions. Bacterial cells with the target concentration of 10^5^ CFU/mL were dispensed into each well. After overnight incubation, wells were inspected visually for bacterial growth. After overnight incubation, 10 μL of each well’s content was transferred to a corresponding well on another microwell plate, containing 150 μL of LB media. This second plate was incubated overnight and the wells were inspected the next day for signs of growth by visual inspection. The minimum concentration of the particles for which no growth was observed was taken as the minimum inhibitory concentration (MIC) value. The tests were also repeated for interaction with different concentrations of AgNO_3_ as a reference material, in the cases of hemin and hemoglobin containing media.

### 2.7 Testing the ability of K_2_EDTA to restore the antimicrobial activity of AgNbO_3_ particles

The test was performed in two formats: solid phase and liquid phase. In the solid phase format, a K_2_EDTA (Becton Dickinson) solution having a concentration of 8 mg/mL was prepared. Different amounts of this solution were spread on 3 Chocolate agar plates, which respectively gave 1600, 800 and 400 µg of K_2_-EDTA per plate surface. After air drying, a bacterial cell suspension (1/100 dilution in TSB of a 0.5 McFarland) was spread over the agar surface using a swab immersed in the cell suspension tube. Then 5 µL of AgNbO_3_ suspensions (50-1600 µg/ml) were loaded onto their designated location on the plate. For a positive control, 5µL of water was dispensed at the center of the plate. Plates were incubated at 35 °C overnight and photographed the next day.

In the liquid phase format, the antimicrobial activity of AgNbO_3_ particles was assessed by microdilution antimicrobial susceptibility testing in six different media, including MHB (Becton Dickinson), with and without 400 µg/mL of K_2_EDTA, MHB with 5 % lysed horse blood (Muller Hinton Fastidious Broth) purchased from Liofilchem (with and without 400 µg/mL of K_2_EDTA), and a filter-sterilized MHB containing Hb at a final concentration of 6 mg/mL (with and without 400 µg/mL of K_2_EDTA). For each medium, each well of a row in a microwell plate was supplemented with different concentrations of AgNbO_3_ particles, in serial one to two dilutions. Bacterial cells with the target concentration of 10^5^ CFU/mL were dispensed into each well. The test was performed on three replicate rows. The plate was incubated overnight and was visually inspected the next day for signs of growth. The minimum concentration of the particles for which no growth was observed was taken as the minimum inhibitory concentration (MIC) value.

## 3. Results and discussion

### 3.1 Antimicrobial activity of antimicrobial AgNbO_3_ particles in the presence of interfering material

The possible impact of blood on the antimicrobial activity of AgNbO_3_ particles is the main focus of the present work as most prospective clinical applications are expected to occur in the presence of blood and blood. The easily replicated approach to quantify this issue is to use three standard plates, one containing no blood (MHA), one containing whole blood (BAP), in which cells are presumed to be intact, and chocolate agar plate (CAP) which contains hemoglobin. The surface minimum inhibitory concentration (sMIC), which in our recent work was shown to be a relevant indicator for applications such as antimicrobial bone-cement [19], was selected as the appropriate quantifier of bacterial toxicity. To measure sMIC, aliquots of serial one to two dilutions of AgNbO_3_ particles are dispensed at delineated spots and are allowed to settle. Then, microbial cells with a nominal count of ∼100 are dispensed at each spot. This number is neither too low as to raise the possibility of sampling error, nor too high as to involve overcrowding and make it impossible to discriminate between circles with no inhibition and those intermediate cases when a fraction of cells survived the antimicrobial action and formed colonies.

The results of bacterial growth on three different agar plates, in the presence of varying concentrations of AgNbO_3_ the particles are shown in Fig 1. We observed that in the case of MHA plates, which lacks any blood product, almost no bacterial cells survived at the surface concentration of 10 ng/mm^2^, and considered this concentration as the surface MIC (sMIC). For BAP plates, having presumably non-lysed red blood cells, the sMIC increased reproducibly from 10 µg/mm^2^ to 20 µg/mm^2^. In the case of CAP plates, having hemoglobin as part of in their composition, the sMIC increased to over 80 µg/mm^2^, an 8× fold increase as compared to the MHA plate.

**Fig 1.**
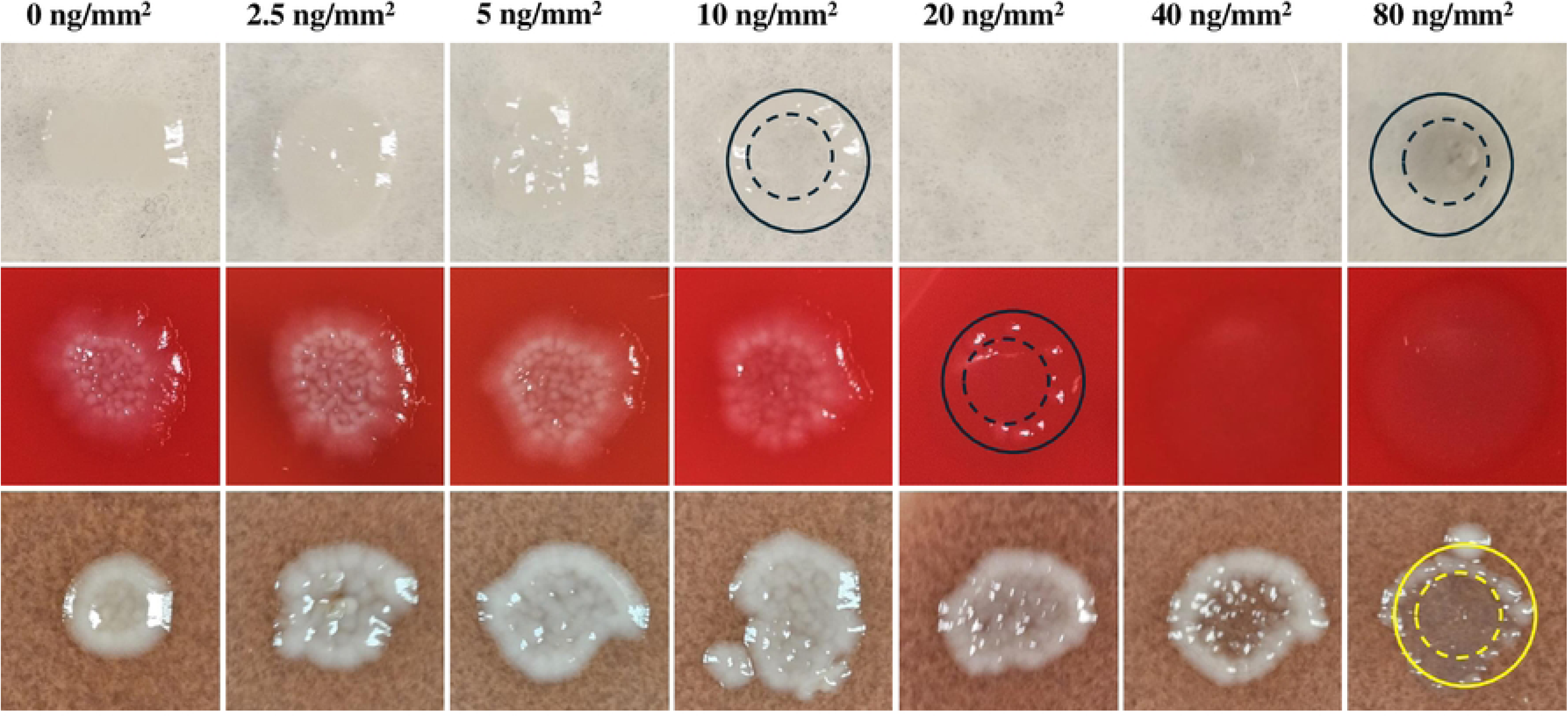
Photographs of the spots of bacterial growth on plates where the particles had been dispensed and which then received a bacterial load of ∼100 CFU. From top to bottom, respectively, MHA, BAP, and CAP. The concentric dashed and solid circles drawn on the representative cases, respectively, indicate the regions of plateaued and low AgNbO_3_ particle concentrations (see below).

Some comments related to the bacterial colony distribution over the spots, particularly those containing AgNbO_3_ concentrations close to the sMIC value, are warranted. For instance, in the case of the CAP plate, at both 40 µg/mm^2^ and 80 µg/mm^2^ the growth in the central region is partially inhibited, while there is a circumpherential ring of higher growth. This is likely the result of the evaporation of sessile droplets containing non-volatile solutes, resulting in the formation of ring-like residues along the contact line pinning [20]. AgNbO_3_ particles are expected to settle down without significant outward radial displacement due to their high mass density of 6.8 g/cm^3^ [21], thus resulting in a post-drying particle distribution that is a projection of the particle distribution in a flattened drop on the hydrophilic plate. Referring to Fig 2, this distribution has a nearly uniform profile in the central regions, as is actually observed by the darker color of the high intensity spot on MHA (inside the dashed circle at the upper right corner in Fig 1), and reduced surface concentration in the region between the dashed and the solid circles. In contrast, the bacterial cells, due to their small mass density, will relocate preferentially to the peripheral region (between the dashed and solid circles), where the particles are less numerous than in the center region. Therefore, the cells finally settled in the central zone are in contact with more particles and feel their antimicrobial impact to a greater extent.

**Fig 2:**
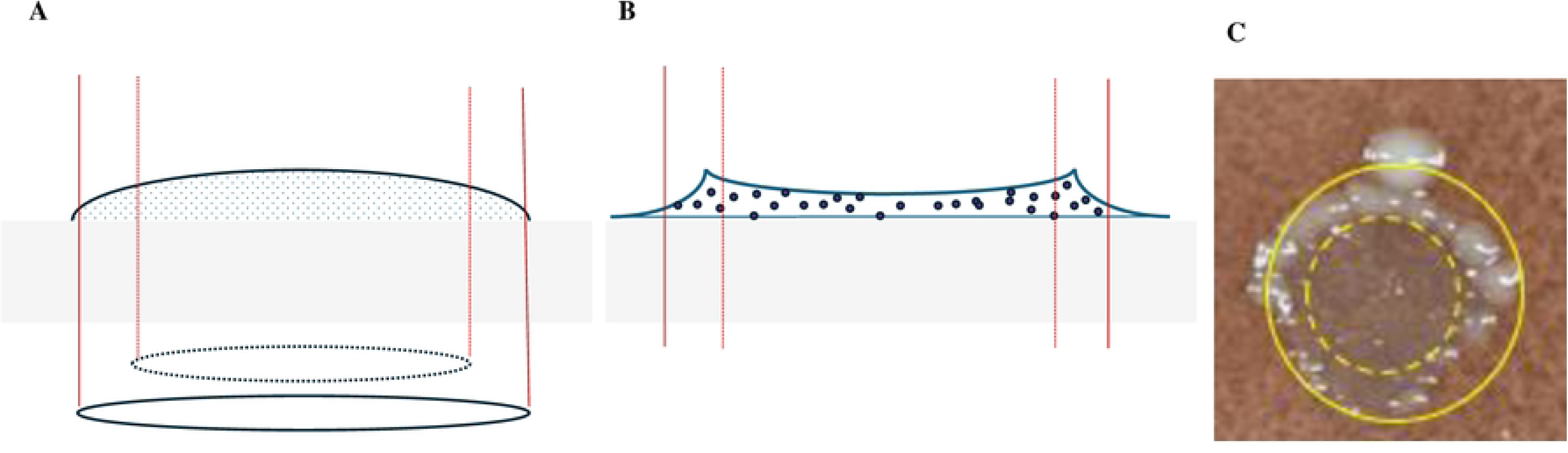
Schematic representation of the distribution at the late drying stage of AgNbO_3_ particles (A) and microbial cells (B) on a hydrophobic gel surface. The two concentric circles, respectively, correspond to the two circles that we had overlaid on the spots of Fig 1, an example of which is presented in (C).

We also studied the dependence of sMIC on the plate composition by a complementary method, where a bacterial cell suspension was spread over the whole area of the three types of plates and left to air-dry. Then, 5 µL of AgNbO_3_ suspension, with concentrations varying by factor of 2-fold in the 50-1600 µg/mL range were dispensed at designated locations on the plates. These drops would give rise to the surface concentrations of 5-160 ng/mm^2^ after drying. The post-incubation photos are presented in Fig 3. Clearly, while the MIC of the AgNbO_3_ particles on BAP was about 2× higher than on MHA, which could be attributed to a small amount of red blood cell lysis in that medium, there is a large increase in the MIC on CAP. The results agree with the sMIC determined via the previous approach. This is supporting evidence for the inhibition of AgNbO_3_ particles’ antimicrobial activity on plates containing either lysed blood or hemoglobin.

**Fig 3.**
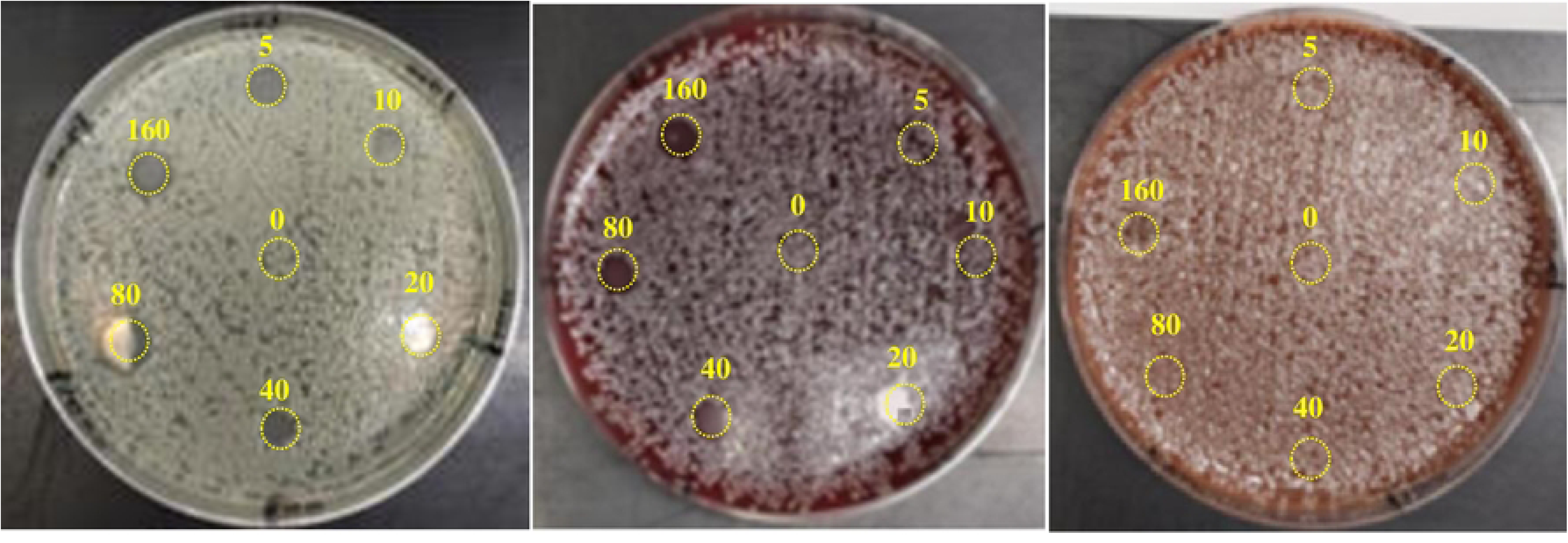
Antimicrobial activity of AgNbO_3_ particles on different plates. From left to right, respectively, MHA, BAP, and CAP. The indicated numbers corresponding to antimicrobial spots indicate particle concentration in ng/mm^2^. Note that the white spots in 20 and 80 ng/mm^2^ cases of MHA and 20 ng/mm^2^ cases of BAP have full inhibition and the white color is due to glare.

A similar shift in the MIC on BAP and CAP plates was also observed when testing for the antimicrobial activity of AgNO_3_ salt, whose mode of action is attributed to silver ions [22]. The test result, which is presented in Fig 4, apart from illustrating clear increase in the MIC for the cases of BAP and, in particular CAP, also indicates a characteristic feature: We observed an absence of a zone of inhibition beyond the area in which the AgNO_3_ solution was dispensed. Apparently, this observation results from a much slower diffusion of silver ions in a blood containing gel or alternatively, and more likely, to the sequestration of diffusing silver ions, resulting from dissociation of AgNO_3_, by blood [23].

**Fig 4.**
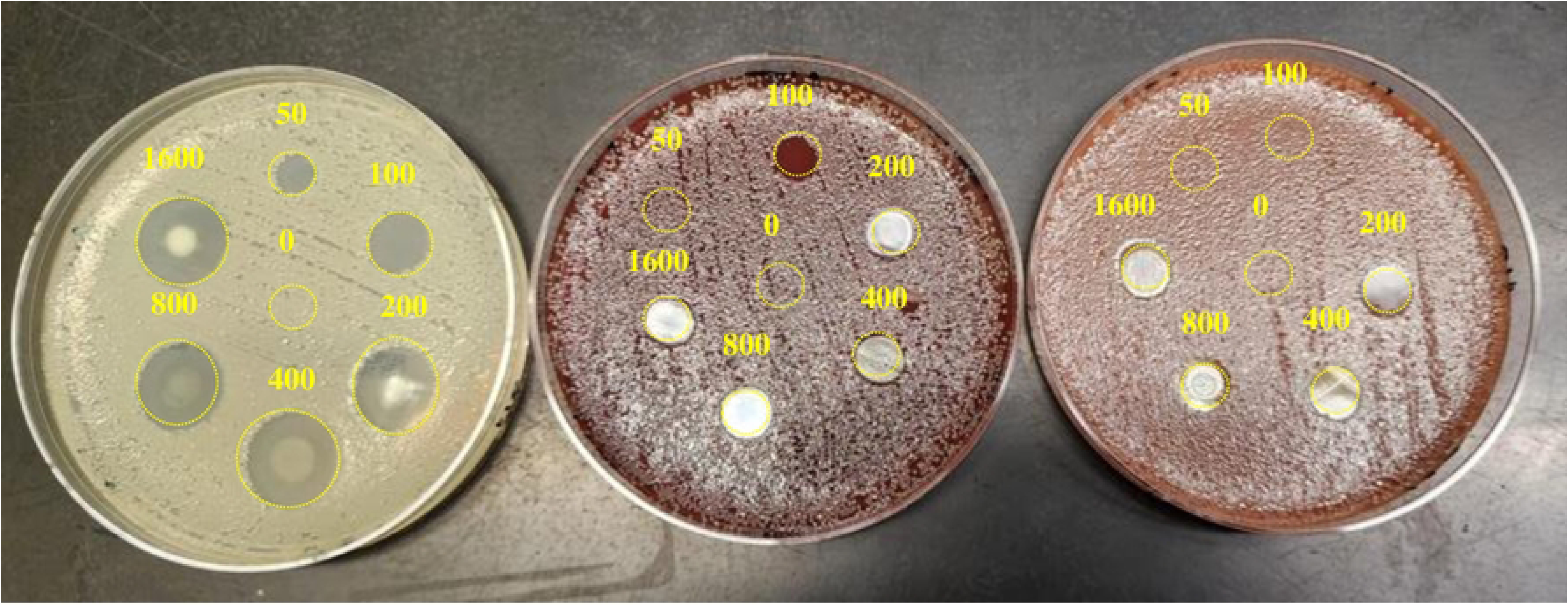
Antimicrobial activity of AgNO_3_ particles on different plates. From left to right, respectively, MHA, BAP, and CAP. The numbers adjacent to the spots indicate the AgNO_3_ concentration in µg/mL. Note that the area of bacterial growth inhibition is diffuse in the case of MHA whereas the diffusion process appears restricted on BAP and CAP. White spots have full inhibition and the white color is due to glare.

The clear shift in the sMIC of the particles on BAP and CAP with respect to the case of MHA may be explained by difference in biochemicals between MHA and BAP/CAP. Both BAP and CAP plates contain hemoglobin. In BAP, red blood cells (RBC) are the source of hemoglobin while in CAP hemoglobin is one of the main components. The partial inhibition by BAP is likely due to the presence of slight amounts of free hemoglobin resulting from ruptured/lysed RBCs in the medium. The attachment of silver particles onto hemoglobin (Hb) occurs at a time and concentration dependent rate as reported by Mahato et al [24]. We therefore investigated whether Hb and hemin were likely causes for the reduced antimicrobial activity of AgNbO_3_ particles.

We first determined the minimum inhibitory concentration (MIC) of the batch of AgNbO_3_ particles used throughout the present work, by assessing its effect on the growth of *Escherichia coli* cells in the presence of the particles suspended in to the microwells at different concentrations as described in the method section. The result of one test replicate is presented in Fig 5 and as it can be seen, the MIC is somewhere in the 10-20 µg/mL range. Over three runs with two replicates each, the MIC fluctuated between 10 and 20 µg/mL.

**Fig 5.**
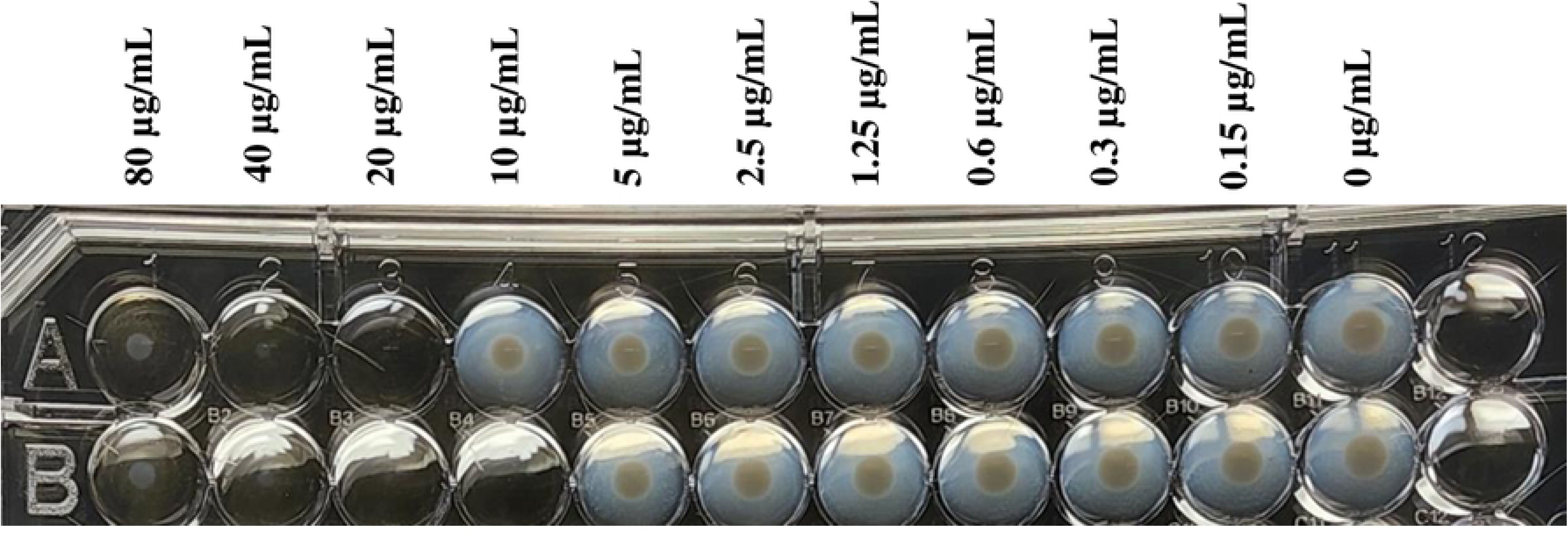
A photograph of the microwell plate used for microdilution test after overnight incubation.

Visual detection of growth and determination of MIC in the presence of interfering compounds, such as Hb, can be difficult. So, subculture from wells onto liquid culture medium was used to confirm the result of MIC determination by visual reading. We observed a significant increase in the MIC, corresponding to loss of antimicrobial activity of the particles, with increasing concentration of both Hb and hemin (Table 1). The inhibitory effect of Hb was found to be higher than that of hemin. This difference is even more apparent when using the molar concentrations of the compounds: Each Hb molecule reduces the antimicrobial activity ∼500 times more than a molecule of hemin. As there are only four times more heme moiety on Hb compared to hemin, we concluded that the heme may not be responsible for the inhibitory effect by Hb. It should be stressed that the reduction in the antimicrobial activity by Hb is not specific to AgNbO_3_ particles, rather it is generic to silver. Indeed, when carrying the test with the AgNO_3_ salt in the presence of Hb, we similarly noted an increase in the AgNO3 MIC against *Escherichia coli*. These observations, combined, demonstrate that free Hb antagonizes the antimicrobial activity of silver compounds with a mechanism where the heme moiety is not the determinant factor.

**Table 1.** The MIC values of AgNbO_3_ particles and AgNO_3_ against *Escherichia coli* for different concentrations of hemoglobin and hemin. To convert to molar concentrations, we have used the molar masses of 651.94 g/mol and 65000 g/mol, respectively, for hemin and hemoglobin.

| Hemoglobin |  |  |  | Hemin |  |  |  |
| --- | --- | --- | --- | --- | --- | --- | --- |
| Con.<br>(mg/mL) | Con.<br>(μM) | MIC<br>AgNbO<sub>3</sub><br>(μg/mL) | MIC<br>AgNO<sub>3</sub><br>(μg/mL) | Con.<br>(mg/mL) | Con.<br>(μM) | MIC<br>AgNbO<sub>3</sub><br>(μg/mL) | MIC<br>AgNO<sub>3</sub><br>(μg/mL) |
| 0 | 0.00 | 16 | 8 | 0 | 0.00 | 16 | 8 |
| 1.25 | 1.92 | 32 | 256 | 1.25 | 0.02 | 32 | 32 |
| 2.50 | 3.83 | 64 | 256 | 2.50 | 0.04 | 32 | 32 |
| 5.00 | 7.67 | 128 | 256 | 5.0 | 0.08 | 32 | 32 |
| 10.0 | 15.34 | 128 | 128 | 10.0 | 0.15 | 64 | 32 |
| 20.0 | 30.68 | 256 | 256 | 20.0 | 0.31 | 64 | 32 |

The use of AgNbO_3_ particles in the presence of Hb released from lysed red blood cells is likely to fail when treating bacterial infections. Therefore, mitigation strategies, ideally involving safe additives to the particles should be investigated. We began with iron due to the ease of inclusion of iron on the particles and on promising findings in the literature, reporting the enhancing effect of iron on conventional antimicrobials [25].

### 3.2 The impact of iron on the antimicrobial activity of AgNbO_3_ particles

The effect of iron ions on the viability of bacterial cells has been studied extensively. Iron, in particular the soluble ferrous form Fe^2+^, is a good electron donor and thus well suited and used as an essential element of the respiratory chain of bacteria cells [26]. Pathogenic bacteria in particular have developed sophisticated systems to acquire sufficient levels of iron as they need iron for the expression of virulence factors and for survival in the host system where ferrous iron (Fe^2+^) is seldom available. The host system, which is for the most part aerobic, allows the presence of the oxidized ferric form Fe^3+^. Even then, iron is not freely available, as, in humans for example, two thirds of iron is sequestered into erythrocytes as heme bound to the oxygen carrying protein hemoglobin [27].

Pathogenic bacteria acquire iron from the host using two mechanisms; The first involves direct contact between the bacterium and the source of iron and the subsequent removal of the iron by its reduction and uptake. The second and the more common, is the synthesis of a high-affinity compound that chelates iron from the source of the iron [28]. Such compounds are named siderophores, of which there are 500 types and without exception all of them have a higher affinity for Fe^3+^ [29]. Siderophore-iron complexes or other iron-binding compounds may then enter the bacterium through specialized high-affinity receptors [30]. Once inside the cell cytoplasm, the iron is reduced to Fe^2+^ and spontaneously released. Elevated levels of Fe^2+^, in combination with H_2_O_2_ (A byproduct of aerobic metabolism or of interaction with silver compounds) can contribute to oxidative stress via the production of reactive oxygen species (ROS) in the form of radical hydroxides through Fenton reaction [31]:

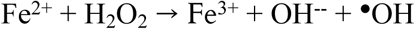

To elucidate the potential mechanism responsible for combinatorial effect of AgNbO_3_ particles and iron ions, we studied the surface charge and elemental composition of the particle surface with more detail, as discussed in the supporting information (see Fig A and Table A of S1 Appendix). We have also demonstrated the generation of ROS in *Escherichia coli* as a result of its interaction with AgNbO_3_ particles using 2’, 7’- dichlorodihydrofluorescein diacetate (H_2_DCFDA) dye which will be described in more detail in a separate submission. The measurement method is presented in the supporting information section (S2 Appendix). Briefly, upon entry of the dye to the live cells, acetate groups of the dye are cleaved by intracellular esterases. Once a nonfluorescent reduced form of dye is oxidized by reactive oxygen intermediates, it is converted to highly fluorescent DCF, whose fluorescence is proportional to the level of intracellular ROS. The test result has shown that the ROS concentrations (As measured by relative fluorescent units or RFU) are at background level for the tested AgNbO_3_ concentrations of 0, 4, and 8 µg/mL. However, at the AgNbO_3_ MIC concentration of 16 µg/mL, the amount of ROS increases from the background level by a statistically significant amount.

We next tested the MIC shifting tests using different concentrations of FeCl_3_ and FeSO_4_ in between 0.1 and 100 µM. FeSO_4_ was selected to test the possible effects of Fe^2+^ according to the Fenton reaction. On the other hand, FeCl_3_ dissociates to Fe^3+^, which isn’t expected to exert influence on the AgNbO_3_ antimicrobial activity as the Fenton reaction is irreversible [32]. The determined MIC in the presence of either FeCl_3_ or FeSO_4_ is presented in Table 2. As it is noted, despite expectations, the presence of Fe^2+^, a reactant for the Fenton reaction, worsened the antimicrobial activity of AgNbO_3_ by two-folds. In contrast, Fe^3+^ improved antimicrobial activity by two-folds. That said, the influence of either iron salts on the AgNbO_3_ MIC may not be as significant as the reported synergism of silver/transition metals of up to 8-fold against certain bacterial species [16].

**Table 2.** The MIC values of AgNbO_3_ particles against *Escherichia coli* at the presence of different concentrations of FeCl_3_ and FeSO_4_.

| $\text{FeCl}_3$ | | $\text{FeSO}_4$ | |
| --- | --- | --- | --- |
| Conc. ($\mu\text{M}$) | MIC ($\mu\text{g/mL}$) | Conc. ($\mu\text{M}$) | MIC ($\mu\text{g/mL}$) |
| 0.1 | 16 | 0.1 | 16 |
| 1 | 8 | 1 | 32 |
| 10 | 8 | 10 | 32 |
| 100 | 8 | 100 | 32 |

Still, we thought that the presence of Fe^3+^ could play a part in the mechanism of antimicrobial action of AgNbO_3_ given that there was enhancement by factor of 2 in the AgNbO_3_ activity when supplying it with Fe^3+^ ions. We should note that iron in the form of Fe^3+^ is already present on the particles due to transfer from balls during the ball milling process as can be noticed from XPS spectra presented and discussed in detail in supplementary data (see Fig B of S1 Appendix). So, we decided to see that if further increasing the iron content of the AgNbO_3_ particles can improve the antimicrobial activity, thereby compensating for the activity loss in the presence of blood products encountered in physiological media. We synthesized AgFe_0.05_Nb_0.95_O_3_ particles and repeated experiments on the antimicrobial activity by the micro broth dilution assay and on the surface of different plates. Unfortunately, there was neither a noticeable change in the value of MIC, nor any observable compensation on the lost activity of the particles on the chocolate agar plate (CAP). Perhaps, the amount of iron on the original AgNbO_3_ particles, transferred via impact with steel balls during its preparation (see Table A of S1 Appendix) has already plateaued the likely enhancement in the antimicrobial activity and adding extra iron on the particles could no longer significantly improve the activity.

### 3.3 Testing the ability of K_2_EDTA in restoring antimicrobial activity of AgNbO_3_ particles

The second approach we adopted for mitigating the antagonizing effect of Hb from lysed red blood cells on the antimicrobial activity of AgNbO_3_ particles was to combine it with a chelating agent. In this regard we took the long known EDTA stimulation seen in hemolysates due to the formation of an iron-EDTA complex [33] as a cue. Interestingly, the published results have illustrated no adverse effect of EDTA on the antimicrobial activity of silver. For instance, Freddi et al [34] have successfully prepared wool–EDTA–Ag complex displaying prominent antimicrobial activity against bacteria.

Another important characteristic of EDTA in the context of our investigation is related to the compound’s anti-biofilm properties, which are synergistic with antimicrobial agents [35, 36]. EDTA acts as a permeating and sensitizing agent for treating biofilm-associated conditions. The structural integrity and resilience of biofilms are attributed to extracellular polymeric substances (EPS), which are a complex mixture of polysaccharides, proteins, nucleic acids and lipids [37]. Cations such as Ca^2+^ and Mg^2+^ act as links between the polysaccharides providing the biofilm its structural rigidity. The chelating effects of EDTA reduces the concentration of these cations in the EPS thereby dissolving the biofilm and enabling direct interactions between the antimicrobial agent and bacteria cells. This mechanism was employed by Said et al. to explain the ability of a wound dressing consisting of silver bound to EDTA and benzethonium chloride capable of three log reduction in viable cells of a biofilm [38]. The result was a great improvement to silver containing dressing without EDTA which caused at most one log reduction of viable bacteria in the model biofilm.

We therefore verified the ability of K_2_EDTA to mitigate the impact of lysed red blood cells on the antimicrobial activity of AgNbO_3_ particles. Results obtained with CAP plates are presented in Fig 6. Different amounts of K_2_EDTA were added prior to spreading bacterial cell suspension. The inclusion of K_2_EDTA at 0.25 µg/mm^2^ reduced the sMIC from >160 ng/mm^2^ to 40 ng/mm^2^, a 4-fold improvement.

**Fig 6.**
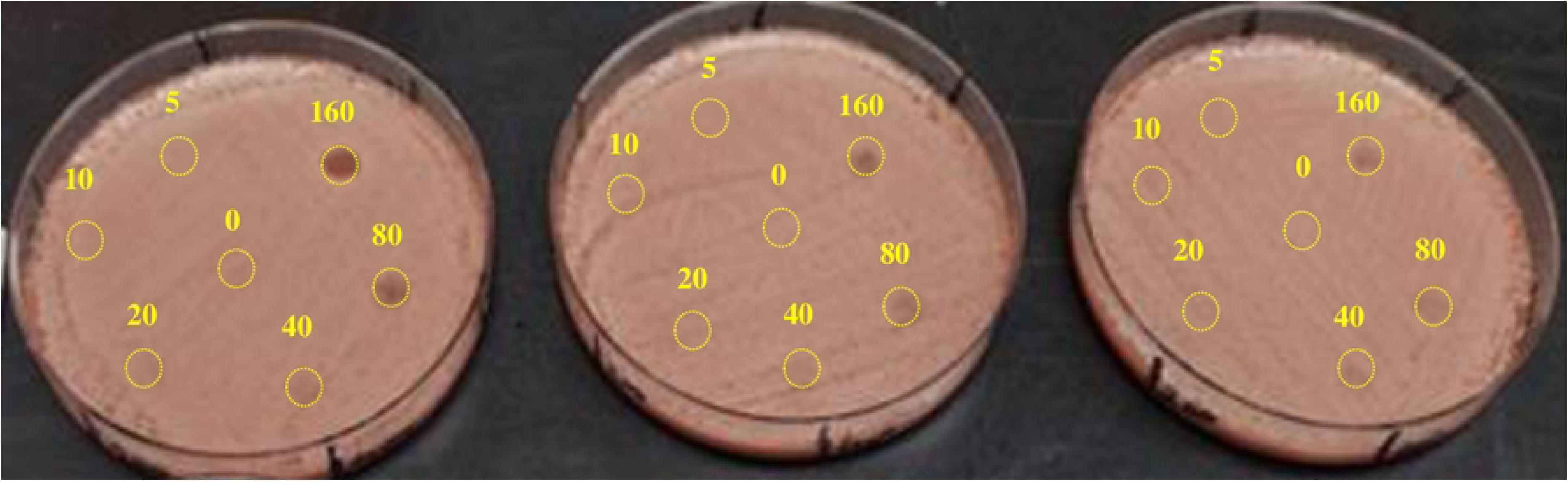
Effect on antimicrobial activity of K_2_EDTA on the antimicrobial activity of AgNbO_3_ particles on CAP plates. The concentrations of AgNbO_3_ particles on each spot are provided in the units of ng/mm^2^. K_2_EDTA amounts on the plates: from left to right: 1600, 800, and 400 µg on the whole surface area of the plate, respectively, corresponding to 0.25, 0.12 5, 0.0625 µg/mm^2^.

The addition of K_2_EDTA also partially restored the inhibitory effect of lysed blood in the micro broth dilution susceptibility format. We utilized three types of broth media: MHB, MHB + %5 lysed blood, and MHB + 6 mg/mL Hb, both in the absence and presence of 400 µg/mL of K_2_EDTA. The main observations are consistent with the results shown in Table 1. In this case, 6 mg/mL of Hb increased the MIC of AgNbO_3_ particles by a factor of 8× while the shift by 5% lysed blood increased the MIC by 16-fold (See Table 3). Adding 400 µg/mL of K_2_EDTA to MHB had no noticeable effect on the MIC when lysed blood or Hb is absent from the media, indicating that K_2_EDTA at this concentration neither inhibits nor augments the antimicrobial activity of the particles. However, the combination of presence of K_2_EDTA with AgNbO_3_ partly neutralizes the inhibitory action of lysed blood and Hb on AgNbO_3_ activity with 4-fold reduction in MICs.

**Table 3.** The MIC values of AgNbO_3_ particles against *Escherichia coli* at the presence of lysed blood and Hb, in the absence and presence of 400 µg/mL of K_2_EDTA.

| | No EDTA | | | With 400 $\mu\text{g/mL}$ of $\text{K}_2\text{EDTA}$ | | |
| --- | --- | --- | --- | --- | --- | --- |
| Media | MHB | MHB 5%<br>Blood | MHB 6<br>mg/mL<br>Hb | MHB | MHB 5%<br>Blood | MHB 6<br>mg/mL<br>Hb |
| MIC<br>($\mu\text{g/mL}$) | 16 | 256 | 128 | 16 | 64 | 32 |

## Conclusion

We demonstrated that while whole blood slightly inhibits the antimicrobial activity of AgNbO_3_ particles, the inhibition by lysed blood is stronger. Accordingly, we decided to test the antimicrobial activity of the particles in the presence of hemoglobin (Hb) and hemin, which result from the lysis of blood cells. While both compounds exhibited strong inhibition, the inhibition per molecule of Hb was over two orders of magnitude greater than that of a molecule of hemin. Therefore, we concluded that the heme moiety, common to the two compounds, was not the main determinant of the inhibition. Attempting to mitigate the inhibitory action of Hb, we tested iron inclusion on the particles to no avail. By contrast, including K_2_EDTA proved successful in countering the inhibitory impact of Hb.

## Supporting information

**S1 Appendix. The iron content of AgNbO3 particles transferred during ball milling**

**S2 Appendix Methodology for illustrating the involvement of reactive oxygen species (ROS) in antimicrobial activity of AgNbO3 particles**

## References

(1) Sim W, Barnard RT, Blaskovich MAT, Ziora ZM. Antimicrobial silver in medicinal and consumer applications: a patent review of the past decade (2007–2017). Antibiotics 2018, 7 (4), 1–12. doi: 10.3390/antibiotics7040093

(2) Meher A, Tandi A, Moharana, Chakroborty S, Mohapatra SS, Mondal A, Dey S, et al. Silver nanoparticle for biomedical applications: A review. Hybrid Advances 2024, 6, 1–12. doi: 10.1016/j.hybadv.2024.100184

(3) Li LJ, Chu C, Yu OY. Application of Zeolites and Zeolitic Imidazolate Frameworks in Dentistry—A Narrative Review. Nanomaterials 2023, 13 (2973), 1–11. doi: 10.3390/nano13222973

(4) Salim MM, Makek NANN. Review of modified Zeolites by surfactant and Silver as antibacterial agents. Journal of Advanced Research in Materials Science 2017, 36 (1), 1–12.

(5) Chen S, Popovich J, Iannuzo N, Haydel SE, Seo D. Silver-ion-exchanged nanostructured zeolite X as antibacterial agent with superior ion release kinetics and efficacy against methicillin-resistant Staphylococcus aureus. ACS applied materials & interfaces 2017, 9 (45), 39271–39282. doi: 10.1021/acsami.7b15001

(6) Abram S, Fromm KM. Handling (nano) silver as antimicrobial agent: therapeutic window, dissolution dynamics, detection methods and molecular interactions. Chemistry–A European Journal 2020, 26 (48), 1–21. doi: 10.1002/chem.202002143

(7) Talebpour C, Fani F, Ouellette M, Salimnia H, Alamdari H. Nondegradable Antimicrobial Silver-Based Perovskite. ACS Sustainable Chem. Eng 2022, 10 (15), 4922–4928. doi: 10.1021/acssuschemeng.1c08181

(8) Esteves GM, Esteves J, Resende M, Mendes L, Azevedo AS. Antimicrobial and Antibiofilm Coating of Dental Implants—Past and New Perspectives. Antibiotics 2022, 11 (235), 1–12. doi: 10.3390/antibiotics11020235

(9) Bistolfi A, Ferracini R, Albanese C, Vernè E, Miola M. PMMA-Based Bone Cements and the Problem of Joint Arthroplasty Infections: Status and New Perspectives. Materials 2019, 12 (4002), 1–12. doi: 10.3390/ma12234002

(10) Simões D Miguel SP, Ribeiro MP, Coutinho P, Mendonça AG, Correia IJ. Recent advances on antimicrobial wound dressing: A review. European Journal of Pharmaceutics and Biopharmaceutics 2018, 127 (2018), 130–139. doi: 10.1016/j.ejpb.2018.02.022

(11) Romanò CL, Tsuchiya H, Morelli I, Battaglia AG, Drago L. Antibacterial coating of implants: are we missing something? Bone & Joint Research 2019, 8 (5), 19–204. doi: 10.1302/2046-3758.85.BJR-2018-0316

(12) Talebpour C, Fani F, Laliberté-Riverin S, Vaidya R, Salimnia H, Alamdari H, et al. Long-Term Prevention of Arthroplasty Infections via Incorporation of Activated AgNbO_3_ Nanoparticles in PMMA Bone Cement. ACS Applied Bio Materials 2024, 7 (6), 4042. doi: 10.1021/acsabm.4c00373

(13) Mulley G, Jenkins ATA, Waterfield NR. Inactivation of the Antibacterial and Cytotoxic Properties of Silver Ions by Biologically Relevant Compounds. PLoS ONE 2014, 9 (4), 1–8. doi: 10.1371/journal.pone.0094409

(14) Gnanadhas DP, Thomas MB, Thomas R, Ralchur AM, Chakravortty D. Interaction of silver nanoparticles with serum proteins affects their antimicrobial activity in vivo. Antimicrobial agents and chemotherapy 2013, 57 (10), 4945–4955. doi: 10.1128/AAC.00152-13

(15) Croes M, de Visser H, Meij BP, Lietart K, van der Wal BCH, Vogely HC, et al. Data on a rat infection model to assess porous titanium implant coatings. Data in brief 2018, 21, 1642–1648. doi: 10.1016/j.dib.2018.10.157

(16) Vasiliev G, Kubo A, Vija H, Kahru A, Bondar D, Karpichev Y, et al. Synergistic antibacterial effect of copper and silver nanoparticles and their mechanism of action. Scientific Reports 2023, 13 (1), 1–13. doi: 10.1038/s41598-023-36460-2

(17) Garza-Cervantes JA, Chávez-Reyes A, Castillo EC, García-Rivas G, Ortega-Rivera OA, Salinas E, et al. Synergistic antimicrobial effects of silver/transition-metal combinatorial treatments. Scientific reports 2017; 7(1), 1–14. doi: 10.1038/s41598-017-01017-7

(18) Raja FNS, Worthington T, Martin R. The antimicrobial efficacy of copper, cobalt, zinc and silver nanoparticles: alone and in combination. Biomedical Materials 2023; 18(4), 1–9. doi: 10.1088/1748-605X/acd03f

(19) Talebpour C, Fani F, Salimnia H, Ouellette M, Alamdari H. Growth dynamics of Escherichia coli cells on a surface having AgNbO_3_ antimicrobial particles. PLOS ONE 2024, 19 (8), 1–12. doi: 10.1371/journal.pone.0305315

(20) Mampallil D, Eral HB. A review on suppression and utilization of the coffee-ring effect. Advances in Colloid and Interface Science 2018, 252, 38–51. doi: 10.1016/j.cis.2017.12.008

(21) Tian Y, Jin L, Zhang H, Xu Z, Wei X, Politova ED, et al. High energy density in silver niobate ceramics. Journal of Materials Chemistry A 2016, 4(44), 17279–17287. doi: 10.1039/C6TA06353E

(22) Mosselhy DA, El-Zaziz MA, Hanna M, Ahmed MA, Husien MM, Feng Q. Comparative synthesis and antimicrobial action of silver nanoparticles and silver nitrate. Journal of Nanoparticle Research 2015, 17, 1–10. doi: 10.1007/s11051-015-3279-8

(23) Walker M, Parsons D. The biological fate of silver ions following the use of silver-containing wound care products–a review. International wound journal 2014, 11 (5), 496–504. doi: 10.1111/j.1742-481X.2012.01115.x

(24) Mahato M, Pal P, Tah B, Ghosh M, Talapatra GB. Study of silver nanoparticle– hemoglobin interaction and composite formation. Colloids and Surfaces B: Biointerfaces 2011, 88, 141–149. doi: 10.1016/j.colsurfb.2011.06.024

(25) Ezraty B, Barras F. The ‘liaisons dangereuses’ between iron and antibiotics. FEMS microbiology reviews 2016, 40 (3), 418–431. doi: 10.1093/femsre/fuw004

(26) Soriano A, Mensa J. Mechanism of action of cefiderocol. Official journal of the Spanish society of chemotherapy 2022, 35 (Suppl 2), 17. doi: 10.37201/req/s02.02.2022

(27) Palmer LD, Skaar EP. Transition Metals and Virulence in Bacteria. Annual review of genetics 2016, 50, 69. doi: 10.1146/annurev-genet-120215-035146

(28) Ratledge C, Dover LG. Iron metabolism in pathogenic bacteria. Annual reviews in microbiology 2000, 54(1), 884.

(29) Hider RC, Kong X. Chemistry and biology of siderophores. Natural product reports 2010, 27(5), 637–655. doi: 10.1039/b906679a

(30) Wandersman C, Delepelaire P. Bacterial iron sources: From siderophores to hemophores. Annual reviews in microbiology 2004, 58, 611–638. doi: 10.1146/annurev.micro.58.030603.123811

(31) Henle ES, Luo Y, Linn S. Fe^2+^, Fe^3^+, and oxygen react with DNA-derived radicals formed during iron-mediated fenton reactions. Biochemistry 1996, 35, 12212.

(32) Pignatello JJ, Oliveros E, MacKay A. Advanced oxidation processes for organic contaminant destruction based on the Fenton reaction related chemistry. Critical reiews in environmental science and technology 2006, 36 (1), 1–84. doi: 10.1080/10643380500326564

(33) Sannes LJ, Hultquist DE. The basis for EDTA-stimulation of methemoglobin reduction in hemolysates of human erythrocytes. Biochemical and biophysical research communications 1979, 91 (4), 1309–1313.

(34) Freddi G, Arai T, Colonna GM, Boschi A, Tsukada M. Binding of metal cations to chemically modified wool and antimicrobial properties of wool-metal complexes. Journal of applied polymer science 2001, 82, 3513–3519.

(35) Finnegan S, Percival SL. EDTA: An antimicrobial and antibiofilm agent for use in wound care. Advances in wound care 2015, 4 (7), 415–420. doi: 10.1089/wound.2014.0577

(36) Gomez NC, Manetsberger J, Benomar N, Abriouel H. Novel combination of nanoparticles and metallo-β-lactamase inhibitor/ antimicrobial-based formulation to combat antibiotic resistant Enterococcus sp. and Pseudomonas sp. strains. International Journal of Biological Macromolecules 2023, 248, 1–12. doi: 10.1016/j.ijbiomac.2023.125982

(37) Prakash B, Veeregowda BM, Krishnappa G. Biofilms: a survival strategy of bacteria. Current science 2003, 85 (9), 1299–1303.

(38) Said J, Walker M, Parsons D, Stapleton P, Beezer AE, Gaisford S. An in vitro test of the efficacy of an anti-biofilm wound dressing. International journal of pharmaceutics 2014, 474 (1-2), 177–181. doi: 10.1016/j.ijpharm.2014.08.034

